# *De novo* genome assembly for an endangered lemur using portable nanopore sequencing in rural Madagascar

**DOI:** 10.1101/2024.05.09.591673

**Authors:** Lindsey Hauff, Noa Elosmie Rasoanaivo, Andriamahery Razafindrakoto, Hajanirina Ravelonjanahary, Patricia C. Wright, Rindra Rakotoarivony, Christina M. Bergey

## Abstract

As one of the most threatened mammalian taxa, lemurs of Madagascar are facing unprecedented anthropogenic pressures. To address conservation imperatives such as this, researchers have increasingly relied on conservation genomics to identify populations of particular concern. However, many of these genomic approaches necessitate high-quality genomes. While the advent of next generation sequencing technologies and the resulting reduction of associated costs have led to the proliferation of genomic data and high-quality reference genomes, global discrepancies in genomic sequencing capabilities often result in biological samples from biodiverse host countries being exported to facilities in the Global North, creating inequalities in access and training within genomic research. Here, we present the first reference genome for the endangered red-fronted brown lemur (*Eulemur rufifrons*) from sequencing efforts conducted entirely within the host country using portable Oxford Nanopore sequencing. Using an archived *E. rufifrons* specimen, we conducted long-read, nanopore sequencing at the Centre ValBio Research Station near Ranomafana National Park, in rural Madagascar, generating over 750 Gb of sequencing data from 10 MinION flow cells. Exclusively using this long-read data, we assembled 2.21 gigabase, 20,330-contig nuclear assembly with an N50 of 98.9 Mb and a 17,108 bp mitogenome. The nuclear assembly had 31x average coverage and was comparable in completeness to other primate reference genomes, with a 95.47% BUSCO completeness score for primate-specific genes. As the first reference genome for *E. rufifrons* and the only annotated genome available for the speciose *Eulemur* genus, this resource will prove vital for conservation genomic studies while our efforts exhibit the potential of this protocol to address research inequalities and build genomic capacity.

## INTRODUCTION

Madagascar’s endemic lemurs are one of most threatened mammalian groups in the world with 98% of recognized species considered threatened with extinction (IUCN, 2023; Schwitzer et al., 2014). Multiple anthropogenic forces have resulted in rapid and severe environmental change on this island nation in the last decades, including significant reductions of forest cover (Vieilledent et al., 2018) on which many of these vulnerable primate species are dependent. Factoring in intensification of climatic events, such as drastic changes in precipitation and amplification of storm severity and frequency, Madagascar’s fragmented forests and the lemur populations residing within them face critical challenges (Weiskopf et al., 2021).

To address biodiversity imperatives such as this, genomic approaches have become crucial tools in aiding conservation management decisions by identifying populations of particular conservation value (Hohenlohe et al., 2021; Supple & Shapiro, 2018). Offering increased precision of estimates compared to analyses that use limited sets of genetic markers (Allendorf et al., 2010; Hohenlohe et al., 2021), genome-based analyses are of particular importance to conservation as they can robustly assess past and present demographic parameters (Abascal et al., 2016; e.g., Palkopoulou et al., 2015), adaptive potential (e.g., Harrisson et al., 2014; Maier et al., 2022), and extinction risk (e.g., Robinson et al., 2022). A limitation to widespread implementation of conservation genomics is the reliance on high-quality reference genomes (Rhie et al., 2021), which can be costly, laborious, and computationally demanding to produce for non-model organisms of conservation concern. However, the “genomics revolution” brought upon by the advent of next-generation, high-throughput sequencing technologies has resulted in a reduction in costs associated with sequencing and an increase in bioinformatic tools available to handle the considerable amount of data generated by such advances (Schiebelhut et al., 2024).

These developments, in conjunction with growing genome consortia initiatives (Darwin Tree of Life Project Consortium, 2022; e.g., Koepfli et al., 2015; Lewin et al., 2022), have led to the proliferation of publicly available reference genomes (Formenti et al., 2022). While genomic resources for IUCN Red List species have grown more than threefold in a span of three years (Brandies et al., 2019; Hogg et al., 2022), there remains global disparities in access to genomic research with sequencing facilities predominantly located in the Global North (Helmy et al., 2016). This disproportionate distribution of sequencing centers often results in biological materials from biodiverse regions exported and researchers from host countries excluded from genomic work, creating inequalities in participation and training opportunities within genomic research (Hoban et al., 2013; Shafer et al., 2014). However, portable DNA sequencers, like the MinION from Oxford Nanopore Technologies (ONT), have the potential to help mitigate such discrepancies by facilitating genomic work in developing countries (Kigen et al., 2023; e.g., Onwuamah et al., 2021; Rivière et al., 2021), reducing barriers to entry by allowing training and participation on associated genomic protocols while fostering more equitable international partnerships.

Portable MinION sequencing has been employed in remote field conditions in Madagascar to great success. In addition to being able to conduct rapid, *in situ* biodiversity assessments to confirm the presence of vulnerable Danfoss’ mouse lemur (*Microcebus danfossi*), one research team was able to use their mobile genetics laboratories to provide a collaborative molecular workshop for local Malagasy students, who previously lacked of access to equipment and hands-on training as major barriers to pursuing such research (Blanco et al., 2020). Usage of portable ONT sequencing has also produced nanopore-only reference assemblies (e.g., Flack et al., 2023; Pozo et al., 2024), highlighting its potential to generate valuable genomic resources while providing underrepresented researchers pivotal access and training to molecular techniques.

Here, we present the first high-quality reference genome of the red-fronted brown lemur (*Eulemur rufifrons*), assembled exclusively from long-read sequencing efforts conducted within Madagascar. Red-fronted brown lemurs are members of the “true lemur” genus (*Eulemur spp*.), one of the most speciose of all of Madagascar’s lemurs, with species distributed in remaining forest fragments across the island (Markolf & Kappeler, 2013). While being primarily frugivorous, *Eulemur spp*. are medium-sized, arboreal primates that display notable dietary flexibility (Johnson, 2007; Overdorff, 1993). *Eulemur spp*. are often considered more resilient to anthropogenic disturbances than larger-bodied lemurs, perhaps due to behavioral plasticity permitting shifts to cathemeral activity in the presence of higher human disturbance (Donati et al., 2016). Despite this perceived hardiness, *Eulemur spp*. are continuing to dwindle, with all 12 species within the genus listed as at least “Vulnerable” to extinction and declining (IUCN, 2023). Conservation of these species are critical, as the health and biodiversity of Madagascar’s tropical forests are intricately woven to the persistence of diverse, frugivorous lemur communities, with many plant species relying nearly exclusively on a specific lemur taxon for seed dispersal and germination (Wright et al., 2011). As at the time of publication there are no other publicly available annotated reference genomes for the *Eulemur* genus. We anticipate our work will be a valuable genomic resource for conservation efforts across Madagascar as well as a proof of concept for how high quality genome assemblies can be generated for such species of conservation concern.

## MATERIALS AND METHODS

### DNA sample and sequencing

#### DNA sample

To conduct whole genome sequencing within Madagascar, we extracted DNA from tissue of an archived *Eulemur rufifrons* specimen that was opportunistically collected within Ranomafana National Park and stored at -20 ºC at the nearby Centre ValBio Research Station. As all sequencing efforts were performed over the course of two time periods, with the pilot study conducted in August 2022 and the bulk of the sequencing conducted in July 2023, the samples and protocols used were adapted over time to account for protocol improvements and unanticipated challenges associated with conducting genomic work in remote regions, as noted below. Skin (totaling 20 mg) was used as the original tissue sample for genomic DNA extractions with Monarch High Molecular Weight (HMW) Extraction Kit for Tissue (New England Biolabs). Spleen tissue (totaling 25 mg) was used for later extractions with the DNeasy Blood and Tissue kit (Qiagen) as a comprehensive dissection of the specimen occurred between sequencing endeavors and the HMW kit was found to be no longer viable, perhaps due to storage conditions in the rainforest. For all extractions conducted, DNA quantity and quality were assessed via a spectrophotometer, NanoDrop 2000 (Thermo Scientific).

#### Library preparation

During the pilot stage of sequencing for this project, 2 µg of HMW DNA was used as input across two library preparations using the Field Sequencing Kit (Oxford Nanopore SQK-LRK001) and following the manufacturer’s protocol. During the second phase of the project, 6 µg of the remaining HMW DNA and 7 µg of the spleen-extracted DNA were used as input for library preparations with the Oxford Nanopore Ligation Sequencing kit SQK-LSK109, split into three and two library preparations for each extraction, respectively. Modifications as described by Flack et al. (2023) were implemented during the latter sequencing effort to prepare additional library material, allowing for multiple reloadings of the same library and flow cell between washes. Prepared libraries were stored at 4 ºC while awaiting sequencing.

#### Sequencing

All genomic sequencing was conducted within Madagascar at the Centre ValBio Research Station on portable MinION sequencers (Mk1C and Mk1B; Oxford Nanopore Technologies). To operate the MinION Mk1B sequencer, an Alienware laptop was used with the following specifications: Intel Core i7 (14-Core) CPU, NVIDIA GeForce RTX GPU, 64GB of memory. No additional hardware was needed to run the standalone Mk1C.

Using ten R9.4.1 flow cells for sequencing, we used the integrated MinKNOW software (v. 23.04.06) and Guppy basecalling (v. 6.5.7) to conduct a total of 21 sequencing runs, each set up initially for 72-hours with runs manually ended by researcher upon significant pore depletion. During the second phase of sequencing, nuclease flushes were conducted after the completion of runs using the Flow Cell Wash kit (WSH-004) following manufacturer’s protocol, and additional aliquots of prepared library were loaded to perform additional sequencing runs. This modification allowed for up to four aliquots of 12 ul of DNA to be loaded on a given flow cell, significantly reducing the sequencing costs associated with the project. During all sequencing runs, Guppy FAST algorithm was used for initial real-time basecalling.

### Computational methods

All bioinformatic analyses were performed on the Rutgers Amarel high performance computing (HPC) cluster. All scripts used for bioinformatic analyses are available at https://github.com/lhauff/Eulemur_genome_assembly.git.

#### Rebasecalling

To improve accuracy of reads originally generated in real-time by Guppy FAST basecalling, *post hoc* basecalling was performed on all reads with SUP, a higher accuracy algorithmic model using Dorado (v. 0.3.4). Briefly, this was achieved by first converting MinKNOW generated FAST5 files to POD5 format before using Dorado model r941_min_sup_g507 to reanalyze the files. Sequencing summary statistics for post-Dorado rebasecalled reads were analyzed in nanoq (v.0.10.0) and are shown in Table 1.

**Table 1.** *E. rufifrons* nanopore sequencing summary statistics before and after rebasecalling with SUP algorithm using Dorado (v 0.3.4, model r941_min_sup_g507). Raw read estimates were extracted from MinKNOW generated sequencing reports and post-basecalling read statistics were generated with *Nanoq* with all reads across all sequencing efforts considered.

|  | Raw read estimates with HMW DNA | Raw read estimates with non-HMW DNA | Read statistics after rebasecalling |
| --- | --- | --- | --- |
| Number of reads | 7,068,390 | 29,529,670 | 36,596,792 |
| Number of bases (bp) | 19,539,340,000 | 31,403,000,000 | 58,760,468,380 |
| N50 read length (bp) | 16,800 | 1,520 | 2,577 |
| Longest read (bp) | - | - | 966,438 |
| Mean read length (bp) | - | - | 1,605 |
| Median read length (bp) | - | - | 825 |
| Mean read quality (QV) | - | - | 7.33 |
| Median read quality (QV) | - | - | 13.58 |

#### Initial nuclear genome assembly

We next used a *de novo* assembler explicitly designed to accurately process long reads, Flye (v. 2.9.1; (Kolmogorov et al., 2019)), to generate the initial *Eulemur rufifrons* assembly from the rebasecalled reads. Additional read trimming or correction was not necessary as raw basecalled reads are used as Flye input. We selected *nano-hq* was selected as the read input type to account for the higher accuracy reads, and we estimated the genome size parameter as 3.0 Gb to accurately reflect a typical primate genome size.

#### Polishing

We next performed extensive polishing of this draft assembly with Medaka (v. 1.9.1, Oxford Nanopore Technologies). To improve parallelism of this computationally intensive program, the Medaka program was split into the three discrete steps: *mini_align, medaka consensus*, and *medaka stitch*. Model r941_min_sup_g507 was specified during *medaka consensus*.

#### Contamination removal

To check the assembled contigs for contamination (bacteria, archaea, or virus), we used Kraken2 (v. 2.1.3; (Wood et al., 2019)) with the standard database, which flagged non-primate reads for subsequent removal from the genome.

#### Repeat masking

To facilitate the speed and effectiveness of downstream annotation, we soft-masked repetitive regions of the genome using RepeatMasker (v. 4.1.5; (Smit et al., 2013)), with “Primate” indicated as species.

#### Scaffolding

We then scaffolded the masked assembly to a *Lemur catta* reference genome (GCF_020740605.2_mLemCat1.pri, (Palmada-Flores et al., 2022)), the most complete chromosome-level Lemuridae reference genome available at the time of assembly, using RagTag (v. 2.1.0; (Alonge et al., 2022). To appropriately reflect our long-read data, minimap2 parameters were set to ‘-x map-ont’.

#### Coverage and quality statistics

To initially assess assembly coverage and completeness, we first used QUAST (5.2.0). This program provided information regarding genome statistics such as assembly length, average coverage depth, and NG50.

To further investigate the quality of our reference genome and to assess how it compares to other publicly available, annotated Lemuroidea genomes, we used BUSCO (v. 5.7.0; (Manni et al., 2021) to evaluate genome completeness based on expected gene content using Benchmarking Universal Single-Copy Orthologs (BUSCO) databases. We specified the predictor to Miniprot (v.0.13; (Li, 2023)), a protein-to-genome aligner and considered orthologues in vertebrate- (vertebrata_odb10) and primate- (primates_odb10) specific lineage databases. We did so for our *E. rufifrons* genome in addition to the other annotated Lemuroidea genomes (*Lemur catta*, GCF_020740605.2_mLemCat1.pri, (Palmada-Flores et al., 2022); *Propithecus coquereli*, GCF_000956105.1_Pcoq_1.0, (Guevara et al., 2021); and *Microcebus murinus*, GCF_000165445.2_Mmur_3.0, (Larsen et al., 2017)) for comparative purposes.

To generate the input files necessary to create a Blobtools2 directory for *Eulemur rufifrons* assembly visualization, we used blobtoolskit (v.4.3.5) and the BUSCO generated results.

#### Gene annotation

To facilitate annotation, we generated pairwise alignment chains between our scaffolded *E. rufifrons* assembly and an annotated human reference genome (GRCh38.p14, (Schneider et al., 2017)) following the make_lastz_pipeline (https://github.com/hillerlab/make_lastz_chains) which uses LASTZ (v. 1.04.22). We then used TOGA (v. 1.1.5; (Kirilenko et al., 2023)) for homology-based gene annotation. TOGA (Tool to infer Orthologs from Genome Alignments) projects orthologous loci for each reference transcript present in the provided reference genome, allowing for annotation and classification of the nearly 20,000 genes present in the gene model we used. Isoform and BED files (curated by the TOGA team from the human GENCODE V38 (Ensembl 104) gene annotation) necessary to obtain meaningful results were acquired within the TOGA GitHub (https://github.com/hillerlab/TOGA/TOGAInput/human_hg38) while the FASTA and GFF files for human genome was obtained directly from NCBI database (GCF_000001405.4).

#### Mitochondrial genome

To accurately assembly the mitochondrial genome, all rebasecalled reads were mapped to the *Lemur catta* mitogenome using minimap2 (v. 2.14; (Li, 2021) and aligned reads were subsequently downsampled to 10,000 reads with SeqKit (v. 2.5.1). With these subsampled reads, the mitogenome was assembled using Flye metagenome mode. The resulting circular contig with 273x coverage was recovered and polished using Medaka (v. 1.9.2) as described previously. MITOS2 within the Galaxy web platform (Afgan et al., 2016) was consulted to confirm that the circular genome comprehensively contained mitochondrial genes. QUAST assembly statistics were calculated for the final mitogenome assembly as well.

## Results

### Sequencing and rebasecalling

We extracted DNA from the tissue and spleen of an *Eulemur rufifrons* specimen which had been opportunistically found within Ranomafana National Park and biobanked at the nearby Centre ValBio Research Station. High molecular weight extractions of the tissue sample yielded approximately 15 µg while DNA extractions from the spleen yielded over 25 µg. The sequencing efforts associated with the pilot stage resulted in lower sequencing success than our second attempt. The two distinct runs across two MinION flow cells (R9.4.1) loaded with high molecular weight (HMW) DNA library prepared with Field Sequencing Kit (Oxford Nanopore SQK-LRK001) generated only 6.5 Gb of data and just over 180,000 reads. However, these efforts validated our field-adapted protocol in rural Madagascar and provided training opportunities to Malagasy researchers on all associated genomic techniques.

Due to the modifications to our protocol as described above, the second stage of in-country sequencing produced significantly improved sequencing data yields. During this stage, DNA was sequenced over 20 days using eight MinION flow cells (R9.4.1) across two portable sequencers (Mk1b and Mk1c). HMW DNA was used initially for library preparations with the Ligation Sequencing Kit (Oxford Nanopore SQK-LSK109). However, upon depletion of that extraction and inexplicable failure of the HMW extraction kit potentially due to field storage conditions, non-HMW DNA from the spleen was used for remaining library preparations with the Ligation Kit as well. In total, 743.96 Gb of data and over 36.4 million reads were generated. The second stage of sequencing allowed for additional training of Malagasy graduate students in associated genomic protocols.

Across both sequencing endeavors, Guppy (v6.5.7) FAST was used for real-time basecalling, with reads being rebasecalled with the higher accuracy SUP algorithm using Dorado (0.3.4). Altogether these reads had a N50 of 2,577 (Table 1), which is expected due to significantly more short reads being generated by the use of non-HMW DNA. Indeed, the average N50 across runs using only HMW DNA was 16.8k.

### Genome assembly and curation

The initial assembly produced by Flye and polished with Medaka consisted of a total length of 2,229,741,522 bp in 25,411 contigs. Kraken2 classified 85.18% of these contigs (N=21,646) to be putatively primate, while 14.02% of the contigs (N=3562) remained unclassified, and 0.799% of these contigs (N=203) were definitively classified as contamination and were subsequently removed. After soft-masking, the assembly was then scaffolded to *Lemur catta*, the most complete chromosome-level Lemuridae reference genome available at the time of assembly. As the specimen sampled was female, any contigs that scaffolded to the Y-chromosome in the *Lemur catta* reference genome were also considered erroneous and removed. The resulting final assembly, EulRuf_1.0, was of length 2,213,739,131 bp and consisted of 20,330 contigs.

The mitochondrial genome assembly built by Flye and polished with Medaka produced a final singular circular contig of 17,108 bp with 1,141x coverage and GC content of 39.08%, as determined by QUAST assembly statistics.

QUAST assembly statistics of the nuclear genome, considering contigs greater or equal to 3000 bp (N=6,823, 2.197 Gb), included an N50 of 98,858,652, an L50 of 9, and an average coverage depth of 31x. The largest scaffold was 294,859,627 bp, which suggests that it has captured the entirety of chromosome 1 (286,674,946 in *Lemur catta* reference). GC content was also very similar to that of the *L. catta* reference genome: 41.06% to 41.36%, respectively.

With an N90 over 30.6 Mb, the bulk of the assembly is represented by 22 contigs (Figure 1A). Based on these assembly quality statistics, our *Eulemur rufifrons* genome compares favorably to other publicly available Lemuroidea genomes (Table 2).

**Table 2:** Assembly statistics for Lemuroidea genomes considered for comparative purposes. Our *de novo* assembly for *Eulemur rufifrons*, generated with long-read portable sequencing conducted entirely in the host-country of Madagascar, is bolded.

| Assembly | Size (Gb) | Contigs/scaffolds/<br>chromosomes | Contig<br>N<sub>50</sub> (Mb) | Scaffold<br>L<sub>50</sub> | Depth | Sample<br>type |
| --- | --- | --- | --- | --- | --- | --- |
| *Lemur catta*<br>mLemCat1.pri | 2.246 | 394/184/30 | 102.2 | 9 | 20.8x | spleen |
| *Microcebus murinus*<br>Mmur_3.0 | 2.487 | 50,982/7,677/34 | 198.2 | 10 | 221.6x | liver,<br>kidney |
| *Propithecus coquereli*<br>Pcoq_1.0 | 2.798 | 299,069/22,538/<br>- | 5.6 | 149 | 104.7x | kidney |
| ***E. rufifrons***<br>**EulRuf_1.0** | **2.213** | **20,330/-/32** | **98.9** | **9** | **31x** | **Skin,**<br>**spleen** |

**Figure 1:**
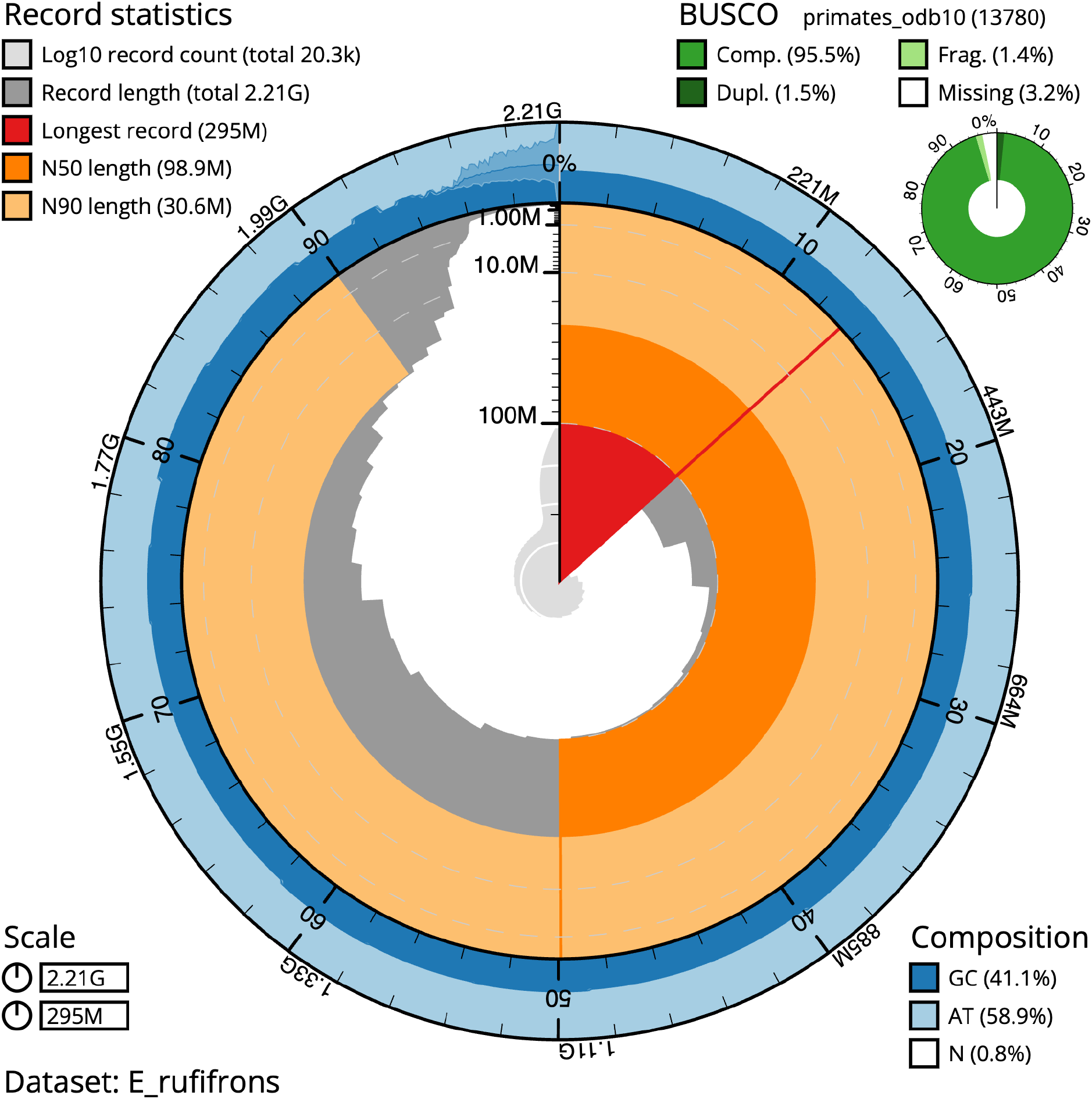
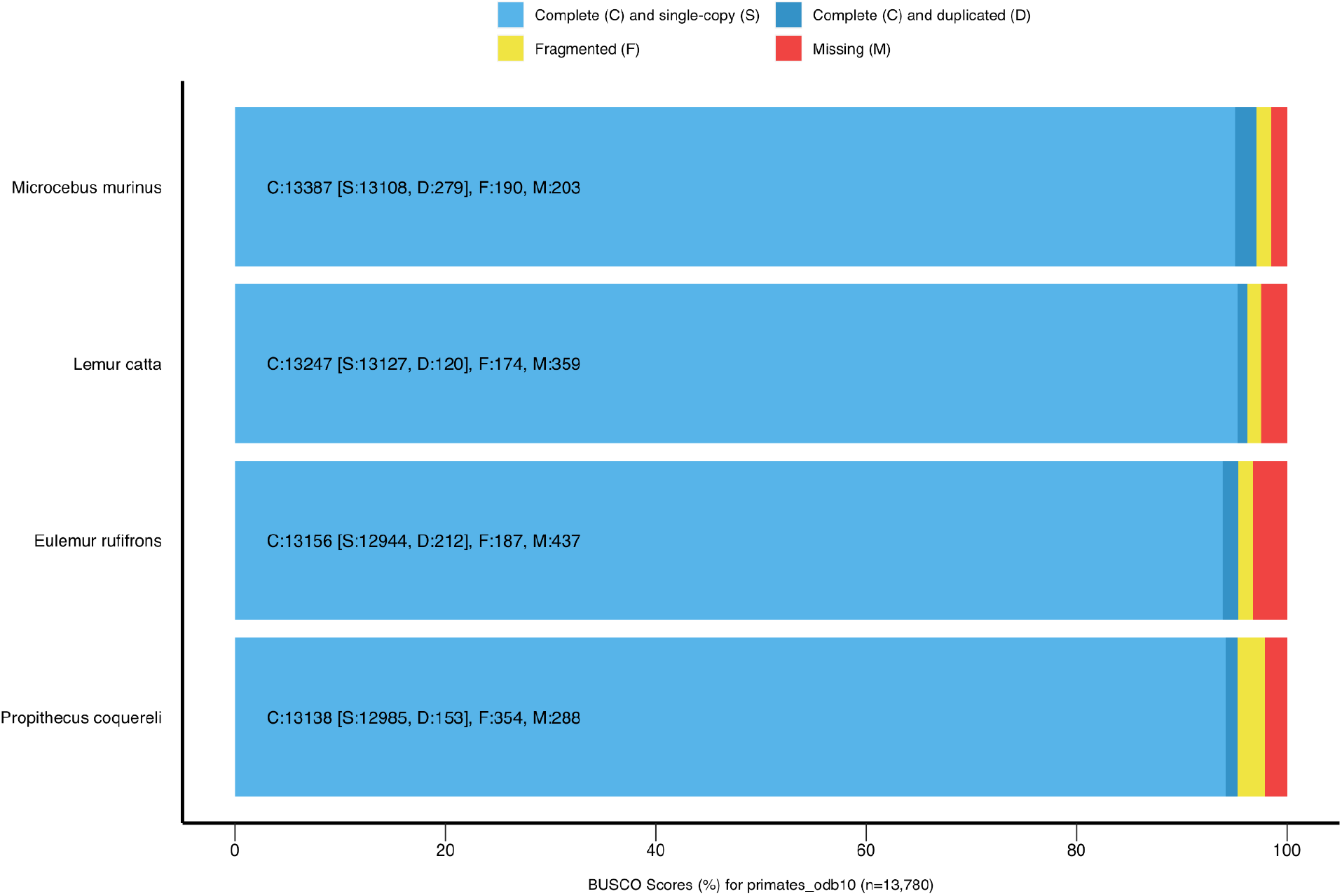
EulRuf_1.0 is a near-complete and highly contiguous genome assembly for red-fronted brown lemur (*Eulemur rufifrons*). (**A)** Final EulRuf_1.0 assembly statistics depicted with BlobtToolKit Snailplot showing total genome size of 2.21 Gb, an N50 of 98.9 Mb, and longest contig over 295 Mb, capturing all of chromosome 1. With an N90 over 30.6 Mb, the bulk of the assembly is represented by 22 contigs. (**B)** Lemuroidea annotated reference assembly completeness comparisons by locally searching primate-specific genes (primates_odb10) with BUSCO (v.5.7.0). With a 95.47% BUSCO completeness score for primate-specific genes, EulRuf_1.0 it is highly comparable to other high-quality, Lemuroidae reference genomes.

**Fig 2:**
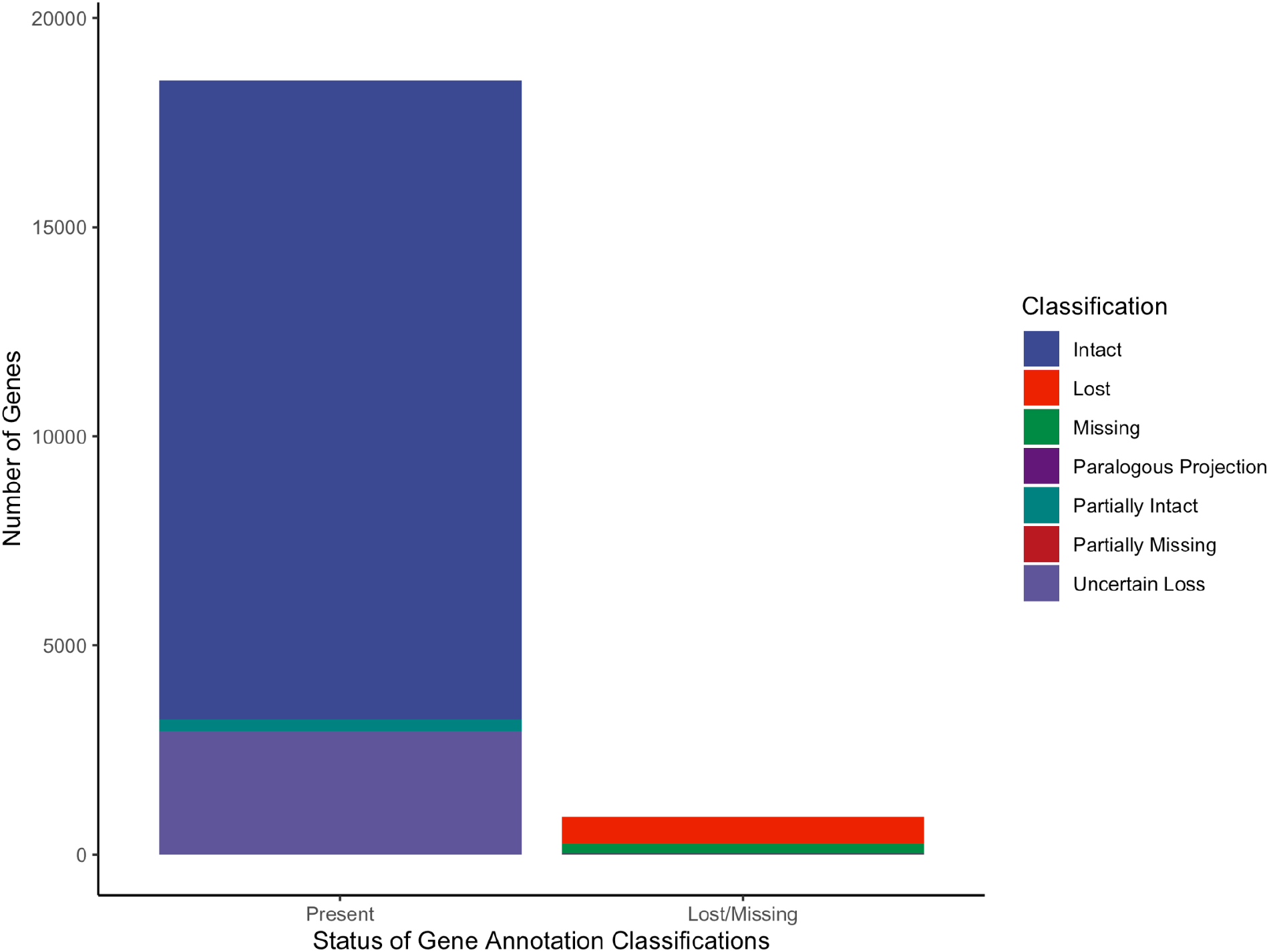
TOGA classifications for 19,414 human reference gene orthologs within *Eulemur rufifrons*. Genes considered to be present with informative annotations include 15,280 “Intact” genes (78.71%), 281 “Partially Intact” genes (1.44%), and 2,947 genes with “Uncertain Loss” (transcripts with a single inactivating mutation in a single exon; 15.18%). Gene annotations that were deemed lost or missing include 634 genes “Lost” (3.26%), 242 genes “Missing” (1.25%), 13 “Paralogous Projections” (0.07%), and 259 genes “Partially Missing” (1.33%).

To calculate comparable annotation completeness scores for all considered lemur genomes (*Lemur catta*, GCF_020740605.2_mLemCat1.pri; *Propithecus coquereli*, GCF_000956105.1_Pcoq_1.0; and *Microcebus murinus*, GCF_000165445.2_Mmur_3.0), we assessed marker gene presence using BUSCO (v.5.7.0; Manni et al., 2021), considering the primate-specific lineage dataset of 13,780 single-copy orthologs (Figure 1B). The EulRuf_1.0 assembly achieved a 95.47% BUSCO completeness score for primate-specific genes, making it highly comparable to other high-quality, annotated Lemuridae reference genomes: *Microcebus murinus* (mmur_3.0) 97.15%, *Lemur catta* (mLemCat1.pri) 96.13%, *Propithecus coquereli* (Pcoq_1.0) 95.34%.

### Genome annotation

We used TOGA to annotate genes in the *E. rufifrons* genome, using the human genome (hg38; GCF_000001405.4) and corresponding isoform and BED files containing 39,664 transcripts of 19,464 genes as a reference (acquired from (https://github.com/hillerlab/TOGA/TOGAInput/human_hg38). Additionally, as TOGA provides classifications regarding exon intactness, which relates to the likelihood that an annotated transcript in the query genome encoding a functional protein (Kirilenko et al., 2023), it can be used to further evaluate genome assembly quality. Of the 19,464 genes initially considered for annotation, TOGA was able to process 19,414, with 232 genes skipped, for various reasons including computational memory requirements. Considering these processed genes, TOGA reported 15,280 genes to be fully intact (78.71%); 281 genes to be partially intact (1.44%), which corresponds to greater or equal to 50% of transcripts considered to be found in coding region; and 634 genes (3.26%) to have at least two inactivating mutations in at least 2 exons, which TOGA considers “lost.” Genes classified as missing more than 50% of transcripts in the coding region due to assembly gaps or fragmentation totaled 259 (1.33%). Transcripts with a single inactivating mutation in a single exon are classified as “uncertain loss” and may be indicative of loss of an exon but not the entire gene (Kirilenko et al., 2023). TOGA classified 2,947 genes (15.18%) as “uncertain loss” in the *E. rufifrons* genome while the remaining 13 genes were deemed “paralogous projections,” indicative of these genes being annotated but with inconclusive projections of transcripts.

With consideration that genes classified as “Intact,” “Partially Intact,” or “Uncertain Loss” by TOGA can all provide useful gene annotation evidence to other annotation tools (Kirilenko et al., 2023), such as EVidenceModeler (Haas et al., 2008), we considered the total of genes with informative annotations to 18,508, or 95.33% of the reference genes assessed. This value is comparable to other genome completeness metrics evaluated for this *de novo* assembly, providing further support that EulRuf_1.0 is a high-quality genome.

## Discussion

Here, we present EulRuf_1.0, the first publicly available, high-quality reference genome for the endangered red-fronted brown lemur (*Eulemur rufifrons*) produced by reads generated entirely within the host-country, Madagascar, on portable sequencers. With a contig N50 of 98.9 Mb and a 95.47% BUSCO completeness score for primate-specific genes, this genome is highly comparable to other annotated Lemuroidea genomes currently available. Annotation efforts with TOGA resulted in 15,280 genes fully annotated, with partial transcripts annotated for an additional 3,228 genes, bringing the total of informative gene annotations to 95.33% of the reference genes assessed. As the only annotated genome available at the time of publication for the speciose and widely distributed *Eulemur* genus, this reference genome will prove to be an important genomic resource for conservation efforts across Madagascar (Thorburn et al., 2023).

In addition to the success of our assembly efforts, these results highlight the potential of using portable sequencing technology to address both the biodiversity crisis and historical underrepresentation of researchers from the Global South in genomics research. With the ability to conduct whole-genome sequencing projects at scale within host countries of high biodiversity, extraction of biological samples to sequencing centers concentrated in the Global North is no longer a genomics imperative. This enables more equitable international partnerships by facilitating enhanced training and participation in genomic work. Although some challenges were encountered while conducting sequencing in a remote region, such as expiration of consumables perhaps due to the tropical environment, our protocol proved to be adaptable and robust. This work joins the growing body of literature that showcase the technological advancements of portable, long-read sequencing technologies and associated bioinformatic processes that enable the creation and curation of reference genomes for species of conservation concern (e.g., Flack et al., 2023; Pozo et al., 2024).

## Acknowledgements

We would like to acknowledge the Ministry for the Environment and Sustainable Development and the Madagascar National Park Service for approving our research permits. For logistical support in the field, the authors want to thank the Centre ValBio (CVB) Research Station staff, especially Dr. Hasina Malalaharivony, and the Madagascar Institute pour la Conservation des Écosystèmes Tropicaux (MICET), especially Dr. Benjamin Andriamihaja. The CVB laboratory is supported by the NSF FSML (1227143) to PCW and its establishment was supported by funding to PCW from IUCN SOS. Additionally, the authors acknowledge the Office of Advanced Research Computing (OARC) at Rutgers, The State University of New Jersey for providing access to the Amarel cluster and associated research computing resources that have contributed to the results reported here. This work is part of the Ph.D. thesis of L. Hauff and was supported by funds granted by the Center for Human Evolutionary Studies, including the Zelnick Award, and by funds awarded by the Rutgers University Ecology and Evolution Graduate Program.

## Data Accessibility Statement

The sequencing data are available via NCBI BioProject: PRJNA1103329. The assembly is available on Dryad: DOI:10.5061/dryad.k3j9kd5gx.

